# SARS-CoV-2 spike protein expressing epithelial cells promotes senescence associated secretory phenotype in endothelial cells and increased inflammatory response

**DOI:** 10.1101/2021.04.16.440215

**Authors:** Keith Meyer, Tapas Patra, Vijay Mahantesh, Ranjit Ray

## Abstract

Increased mortality in COVID-19 often associates with thrombotic and microvascular complications. We have recently shown that SARS-CoV-2 spike protein promotes inflammatory cytokine IL-6/IL-6R induced trans-signaling responses which modulate MCP-1 expression in human endothelial cells. MCP-1 is secreted as a major component of the senescence associated secretory phenotype (SASP). Virus infected or Spike transfected human pulmonary epithelial cells (A549) exhibited an increase in senescence related marker proteins. TMNK; as a representative human endothelial cell line, when exposed to cell culture supernatant derived from A549 cells expressing SARS-CoV-2 spike protein (Spike CM) exhibited a senescence phenotype with enhanced p16, p21, and SA-β-galactosidase expression. Inhibition of IL-6 trans-signaling by Tocilizumab, prior to exposure of supernatant to endothelial cells, inhibited p16 and p21 induction. Likewise, inhibition of receptor signaling by Zanabrutinib or Brd4 function by AZD5153 also led to limited induction of p16 expression. Senescence lead to an enhanced level of adhesion molecule, ICAM-1 and VCAM-1 in human endothelial cells, and TPH1 attachment by *in vitro* assay. Inhibition of senescence or SASP function prevented ICAM/VCAM expression and leukocyte attachment. We also observed an increase in oxidative stress in A549 spike transfected and endothelial cells exposed to Spike CM. ROS generation in TMNK was reduced after treatment with the IL-6 specific inhibitor Tociliximab, and with the specific inhibitors Zanabrutinib and AZD5153. Taken together, we identified that the exposure of human endothelial cells to cell culture supernatant derived from SARS-CoV-2 spike protein expression displayed cellular senescence markers leading to enhanced leukocyte adhesion with coronary blockade potential.

## Introduction

SARS-CoV-2 is the seventh coronavirus known to infect humans. Among these coronaviruses, SARS-CoV, MERS-CoV and SARS-CoV-2 can cause severe disease (1). SARS-CoV and MERS-CoV induce a substantial cytopathic effect and dysregulation of host immune responses. Immune-mediated pathogenesis is likely to be a potential factor for severe outcome in MERS-CoV-infected patients. Infection of mononuclear phagocytes (MNPs) is abortive in SARS-CoV; however, MERS-CoV can replicate in monocytes, macrophages and dendritic cells (2). Evidence of productive SARS-CoV-2 infection in immune cells remains to be determined. Potential mechanisms for disease progression include high rates of viral replication, which could be responsible for enhanced host cell cytolysis, and the strong production of inflammatory cytokines and chemokines by infected epithelial cells perpetuating virtual damage and excessive accumulation of monocytes, macrophages and neutrophils. The disease severity correlates with the number of inflammatory cytokines present in the serum. The role of SARS-CoV-2 induced excessive inflammatory responses as a factor contributing to disease severity as a major outcome needs to be critically examined. Acute kidney injury (AKI), cardiac damage and abdominal pain are the most commonly reported co-morbidities of COVID-19 (3, 4). SARS-CoV-2 infection may associate with damage occurring due to specific pathogenic conditions, including cytokine release syndrome (5).

Cellular senescence is a cell state triggered by stressful insults and certain physiological processes, characterized by a prolonged and generally irreversible cell-cycle arrest with secretory features, macromolecular damage, and altered metabolism (6). Senescent cells secrete a plethora of factors, including pro-inflammatory cytokines and chemokines, growth modulators, angiogenic factors, and matrix metalloproteinases (MMPs), collectively termed the senescent associated secretory phenotype (SASP) (7, 8). The SASP constitutes a hallmark of senescent cells and mediates many of their patho-physiological effects.

SARS-Cov-2 is accompanied by thrombosis, vascular damage and coagulation. These are caused by the release of Von Willebrand factor (VWF). VWF is a glycoprotein and is one of several components of the coagulation system that work together. VWF is released from endothelial cells that line blood vessels and contribute to blood clots during COVID-19 (9). VWF is somewhat unique in expression to endothelial cells. IL-6 induced ULVWF (Ultra long Von Willebrand Factor) release only when it was in complex with the soluble IL-6 receptor. IL-6, but not IL-8 or TNF-α, inhibited the cleavage of ULVWF strings by ADAMTS13 under flowing, but not static, conditions. These results suggest that inflammatory cytokines may stimulate the ULVWF release and inhibit the ULVWF cleavage, resulting in the accumulation of hyperreactive ULVWF in plasma and on the surface of endothelial cells to induce platelet aggregation and adhesion on the vascular endothelium. These findings describe a potential linkage between inflammation and thrombosis that may be of therapeutic importance (10).

The present study addresses critical issues regarding the identification of molecular aspects of viral and host determinants of infection as potential mechanisms for immune-mediated pathology, a highly significant area in COVID-19 research. Our study identified cells - other than lung epithelial cells - supporting SARS-CoV-2 growth, and examined for potential multiorgan pathogenesis. Results from our study suggest that the host determinants and underlying mechanisms of immune-mediated pathology might be predisposing individuals to disease aggravation, and development of potential host directed therapeutic candidates.

## Materials and Methods

### Cell culture and transient transfection

Transformed human lung epithelial cells (A549), liver epithelial cells (Huh7.5), liver sinusoidal endothelial cells (TMNK-1) (kindly provided by A. Soto-Gutierrez, University of Pittsburg, PA), and EAhy926 cells were cultured in Dulbecco’s modified Eagle’s medium (DMEM) (Hyclone) containing 10% fetal bovine serum (FBS) (Sigma), 100 U of penicillin/ml, and 100 mg of streptomycin/ml (Sigma). The cells were maintained in a humidified atmosphere at 37°C with 5% CO_2_.

Cells were sub-cultured in 6-well plate at ∼60% confluency and transfected with plasmid DNA (pcDNA3.1-SARS-Cov-2-Spike MC-0101087-5834, kindly provided from BEI Resources), and empty vector construct (500 ng/plate) using Lipofectamine 3000 (Life Technologies) following the manufacturer’s instruction. Cell lysates were prepared after 72 h of transfection for analyses. Culture supernatant was collected after 72h (A549 Spike CM) in the presence or absence of specific inhibitors.

For infection with SARS-CoV-2, A549 cells were cultured in DMEM containing 2% FBS. SARS-CoV-2 isolate (USA-WA1/2020, BEI Resources) derived from an isolate sourced to Wuhan, China was used to infect cells at a multiplicity of infection of 0.5. All live virus experiments were performed in a P3 facility approved by the Institutional Biosafety Committee. Cell lysates were collected 48 hours after infection.

### Senescence associated β-Galactosidase expression

Initial cellular senescence was determined using the SPiDER-βGal Cellular Senescence Detection kit (Dojindo) Senescence-associated β-Galactosidase in expressed by cells undergoing senescence. Spike transfected A549 were measured at 72 h post-transfection. TMNK cells were measured at 24 h post exposure to A549 Spike conditioned media. Cells were initially exposed to Bafilomycin A1 for 1 hour to inhibit endogenous β-Galactosidase activity. Cells were given SPiDER-βGal reagent and incubated an additional 30 minutes prior to visualization by immunofluorescence following supplier protocol.

### Inhibitor treatment

Tocilizumab (Absolute Antibody), AZD5153 and Zanabrutinib (<u>Selleckehem</u>), ST2825 (Medchem Express), YCG063 (Calbiochem), and AT1 receptor antagonist Candesartan cilexetil (Tocris) were used in this study. The cells were treated with the Candesartan cilexetil (dissolved in DMSO) at 1 µM concentration for 20 h following earlier reports (11, 12). Inhibitors were dissolved in DMSO.

### Immunofluorescence

A549 cells were transfected with SARS-CoV-2 spike protein using an EGFP linker construct by lipofectamine 3000. After 48h, cells were fixed and γH2AX antibody (Cell Signaling) was used for IF. γH2AXwas visualized using anti-rabbit Alexa Fluor 594 (Invitrogen). Nuclei were stained by DAPI.

### Western blot analysis

Cell lysates were electrophoresed to resolve proteins by SDS-PAGE, transferred onto a nitro-cellulose membrane, and blocked with 4% non-fat dry milk. The membrane was incubated at 4°C overnight with specific primary antibody, followed by a secondary antibody conjugated with horseradish peroxidase. The protein bands were detected by chemiluminescence (Life Technologies). The blot from the same run was reprobed with β-actin (Sigma) HRP conjugated antibody to compare protein load in each lane. Commercially available antibodies for phospho-p38 MAPK (Thr180/Tyr182, Thr202/Tyr204), p-AKT (S473), p21, γ-H2AX, VCAM-1, ICAM-1 were procured from Cell Signaling Technologies, and p16 antibody was procured from Santa Cruz Biotechnology.

### Leukocyte adhesion assay

Leukocyte attachment on an endothelial cell surface was performed using monocyte cells (THP-1) with TMNK-1 and EA_hy926_ endothelial cells. Adhesion of THP-1 cells was measured by Cytoselect Leukocyte-Endothelium Adhesion assay kit (Cell Biolabs, Inc) following suppliers protocol.

### ELISA

Cell culture supernatants from SARS-CoV-2 spike protein transfected cells were analyzed for the presence of secreted IL-6, MCP-1, IL-1α (Sigma) and HMGB-1 (Novus Biologicals) using ELISA kits following the manufacturer’s instructions. TMNK-1 cells and from SARS-CoV-2 spike expressing A549 cell culture medium in the presence or absence of inhibitors were used.

### ROS assay

A549 were cultured after transfection of SARS-CoV-2 spike protein following transfection of cells for 48h, and were treated with various inhibitors for 24 hrs. in a 37^0^C CO_2_ incubator. TMNK-1 cells were exposed to A549 spike CM and incubated as above 24h prior to measurement. The intracellular ROS was measured using a commercially available cell based assay detection kit (Cayman Chemicals, MI USA).

### Statistical analysis

Graph Pad Prism 7 was used to analyze the experimental data. All experiments were performed at least three times for reproducibility. The results are presented as mean ± standard deviation. Non-parametric Mann-Whitney U test and paired two-tailed t-test analyses were performed to compare the mean values between the two groups. Statistical significance was considered as *p* <0.05.

## Results

### Ectopic expression of SARS-CoV-2 spike protein in A549 cells induces signs of cellular senescence morphology

Cellular senescence via SASP may play a role in poor COVID-19 patient clinical outcomes (13). Infection of A549 cells *in vitro* with SARS-Cov-2 led to the significant induction of phosphorylated p38 as an associated senescence marker, coupled to an increase in p16 after 48h of infection (Fig.1, panel A). However, there was no apparent induction of phosphorylated Akt or increase in p21 (data not shown). We examined A549 cells transfected with the full-length Spike plasmid and noted that each of these markers were enhanced in cells expressing viral Spike protein (Fig.1, panel B). Further, examination of Spike transfected A549 cells displayed enhanced expression of the senescence marker SA-β-gal by immunofluorescence. The increased expression was seen most prominently in single cells, with lesser fluorescence visible in surrounding cells (Fig.1, panel C).

**FIGURE 1.**
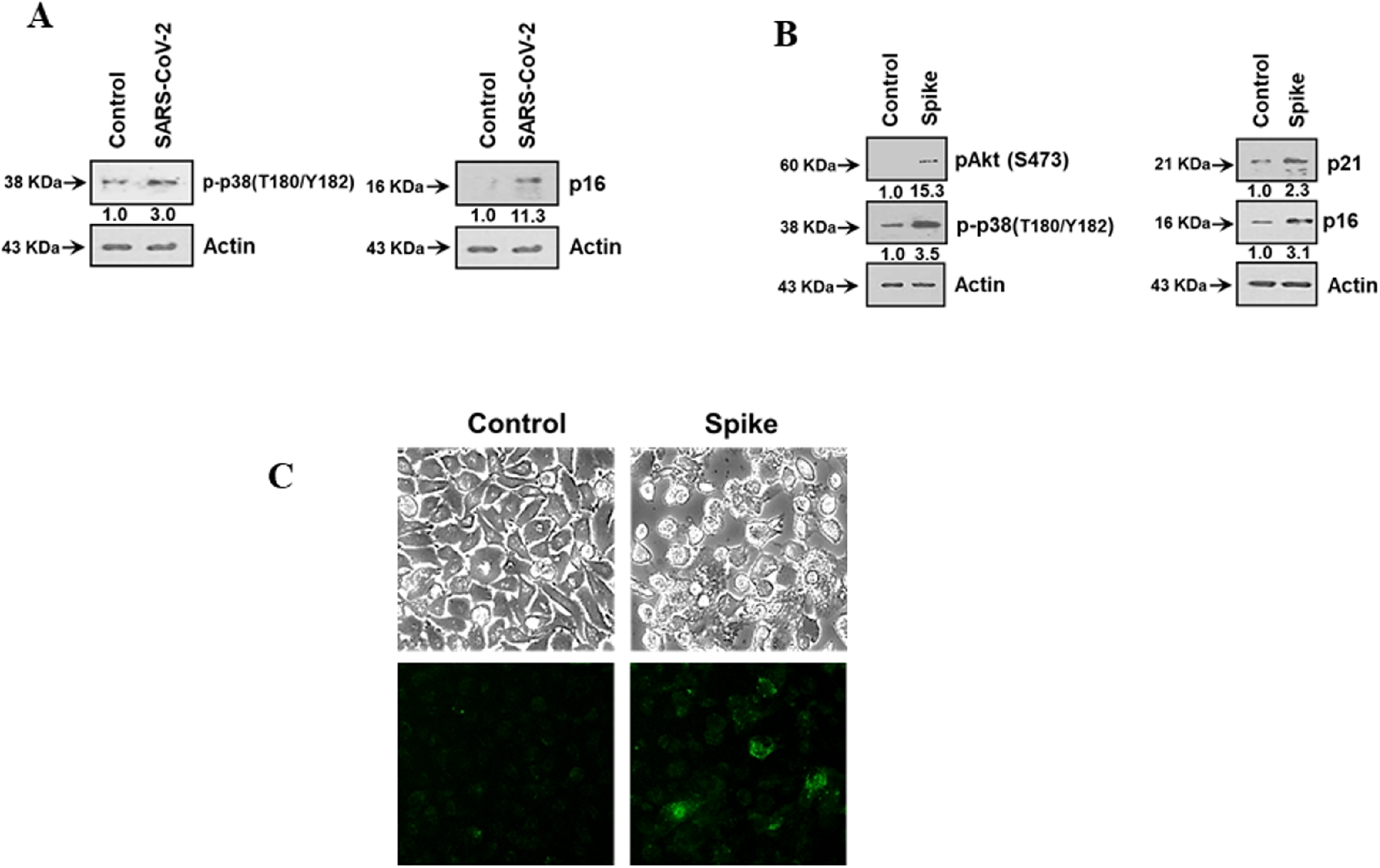
SARS-CoV-2 spike protein expression induced senescence in A549 cells. Cellular expression of SARS-CoV-2 Spike protein exhibits senescence markers and alters cell morphology. SARS-CoV-2 infection of A549 cells displayed senescence associated markers (Panel A). A549 cells transfected with SARS-CoV-2 spike protein induced the expression of the senescence markers p16 and p21, with concurrent increase in expression of phosphorylated Akt and p38 (panel B). Spike transfected A549 also exhibited enhanced SA-β-Gal expression (panel C). Experiments were performed in triplicate.

Oxidative stress has been associated with the pathology of SARS-CoV-2 infection, including its amplification of cytokine storm and coagulopathy (14). ROS generation has been associated with the induction of senescence and maintenance of a viable senescent state in A549 cells was dependent upon ROS (15, 16). Here, we measured intracellular ROS, and observed an approximate 3-fold increase in ROS in A549 Spike transfected cells compared to non-transfected control cells (Fig. 2, panel A). Free radicals are known to cause DNA damage. The increased production of intracellular ROS plausibly contributes to DNA damage leading to cellular senescence. We observed an increase in the DNA damage response (DDR) marker, γ-H2AX, in cells transfected with viral spike (Fig. 2, panel B) and γ-H2AX was found to be localized to the nucleus in A549 cells expressing SARS-CoV-2 Spike protein (Fig. 2, panel C). Brd4 has been observed to be required in senescence immune surveillance and SASP associated paracrine signaling (17). Spike gene transfection of A549 cells led to an enhanced expression of Brd4 (Fig. 2, panel D). Our results indicated that the expression of SARS-CoV-2 Spike protein induces a senescent state in infected or Spike transfected A549 cells, potentially leading to enhanced SASP associated paracrine signaling.

**FIGURE 2.**
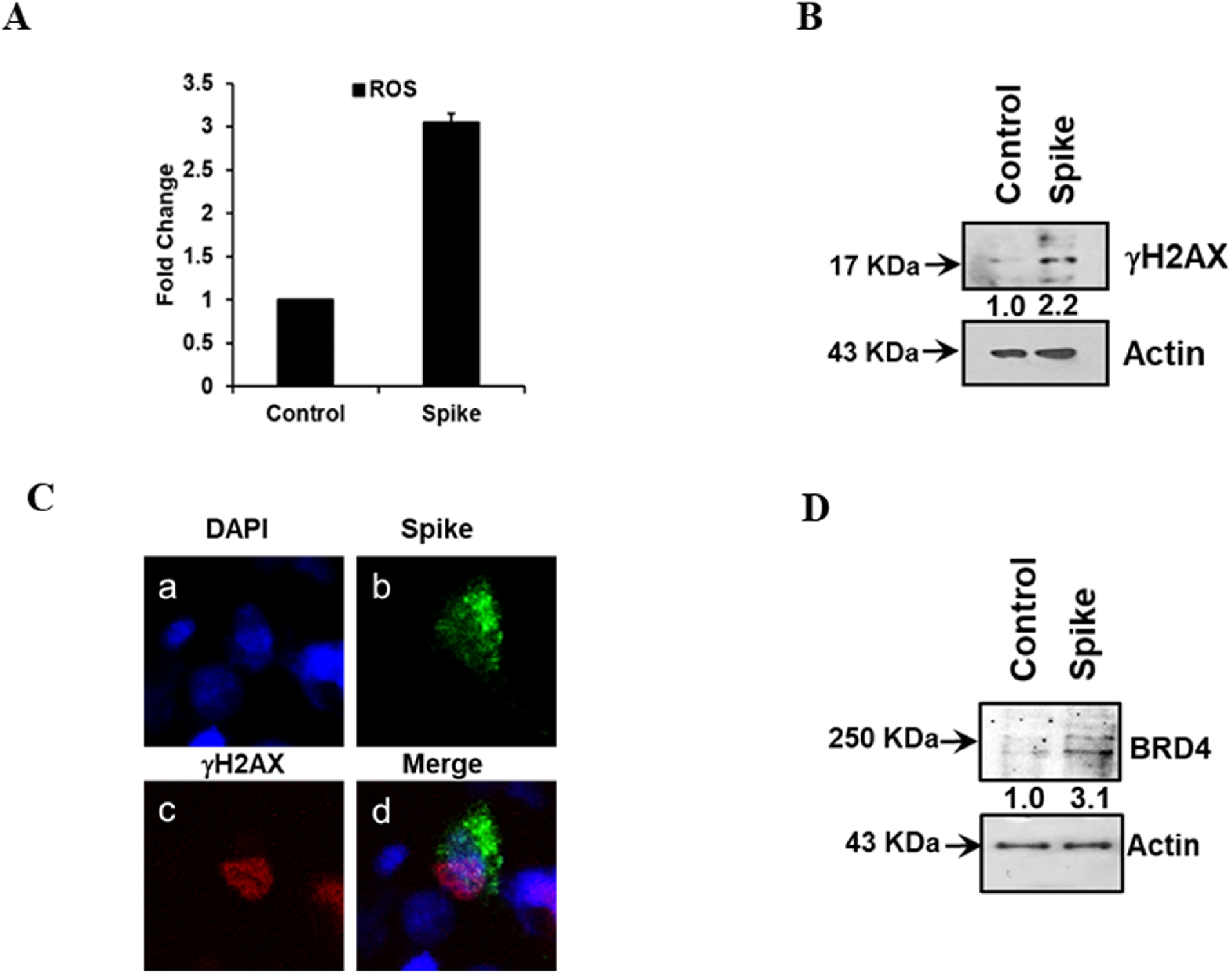
SARS-CoV-2 generates oxidative stress in A549 cells. SARS-CoV-2 Spike transfection induced ROS release (panel A). SARS-CoV-2 Spike expression in A549 cells induced γ-H2AX expression (panel B), nuclear localization of γ-H2AX (panel C), and increased BRD4 expression (panel D). Experiments were performed in triplicate.

### Secretory component from Spike transfected A549 cell culture medium induces senescence in endothelial cells

A549 cells were transfected with the SARS-CoV-2 Spike protein, and incubated for 48 h prior to exposure to the inhibitors Tociliximab, Zanubrutinib, AZD5153 or ST2825 overnight. Spike transfection induced an increase in both p16 and p21 expression in A549 cells. The senescence markers were diminished by the addition of Tociliximab, Zanubrutinib, and AZD5153.

Expression remained unchanged when the cells were exposed to the MyD88 specific inhibitor; ST2825 (Figure 3, panel A). Spike transfected A549 cells exhibited enhanced secretion of alarmins HMGB1, IL-1α, and IL-6 (Fig. 3, panel B). The use of inhibitors decreased cytokine and alarmin release.

**FIGURE 3.**
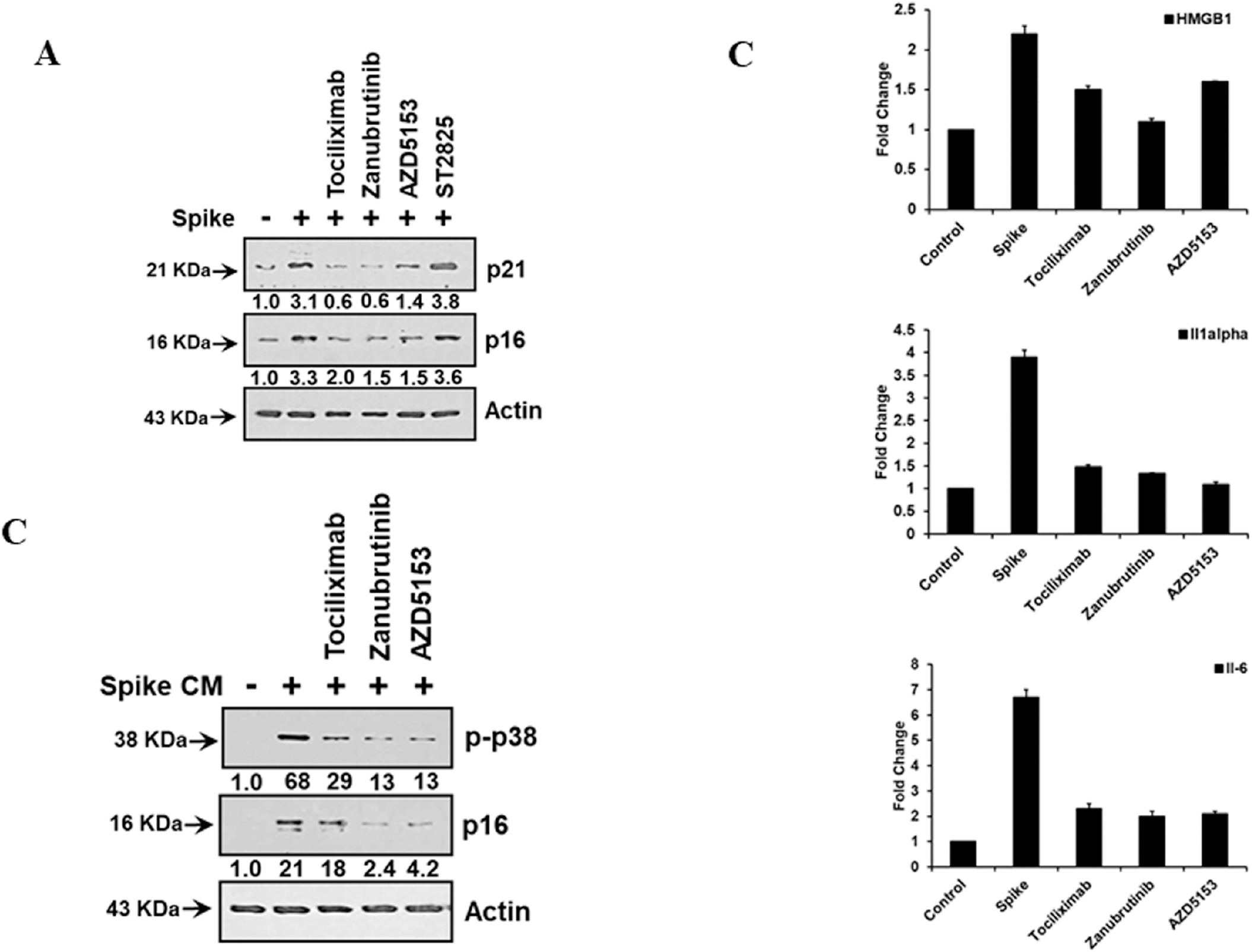
SARS-CoV-2 Spike transfected A549 display senescence markers and enhanced cytokine release. Spike transfection induced an increase in both p16 and p21 expression and were inhibited by the addition of Tociliximab, Zanubrutinib, and AZD5153 inhibitors (panel A). Spike transfected A549 cells exhibited enhanced secretion of alarmins (HMGB1 and IL-1α) and IL-6 (panel B). TMNK-1 cells were incubated with CM generated in the presence of specific inhibitors as above, and the level of the senescence markers were reduced (panel C). Experiments were performed in triplicate.

Culture media (CM) derived from these treated cells were added to the endothelial cell line TMNK, and incubated overnight. Cell lysates derived from A549 Spike CM treated TMNK cells exhibited enhanced levels of phosphorylated p38 as an associated senescence marker, and increases in p16. When these A549 Spike CM were generated; as above, in the presence of specific inhibitors, the levels of the senescence markers p16 and p21 were reduced by treatment of transfected A549 cells with Zanubrutinib or AZD5153 (Fig. 3, panel C).

Endothelial cells are the source of pro-inflammatory cytokine release during infection (18). We examined whether the culture supernatant from spike transfected A549 cells induce cellular changes in TMNK. As we observed an enhancement of SA-β-Gal and major senescence markers in A549 transfected with SARS-CoV-2 Spike expression, we analyzed the endothelial cell line;

TMNK, post exposure to culture medium from these transfected A549 cells. Microscopically, there was a marked reduction in cell number over time, with cells appearing with a characteristically flattened shape and a slightly increased size at day 1 post-exposure; and cells expressed the senescence marker, SA-β-gal (Fig.4, panel A). On day 3, the TMNK cell number was greatly reduced in comparison to TMNK exposed to control media, and the visible signs of SA-β-gal expression were enhanced in all remaining cells (data not shown). Myeloid differentiation 88 BTK associated signaling mediate cytokine associated regulation of NF-kB.

MyD88 has been identified as a mediator of epithelial cell senescence derived from TLR/IL-1R mediated signaling (19). Bruton tyrosin kinase (BTK) activates p53 mediated senescence, and inhibition of BTK had a concomitant decrease in senescence markers (20). Exposure of TMNK cells to A549 Spike expressing culture medium led to a marked induction of BrD4, which could be inhibited utilizing Zanabrutinib (Fig. 4, panel B). Therefore, inhibition of upstream pathways associated with senescence may be important to ameliorate SASP establishment.

**FIGURE 4.**
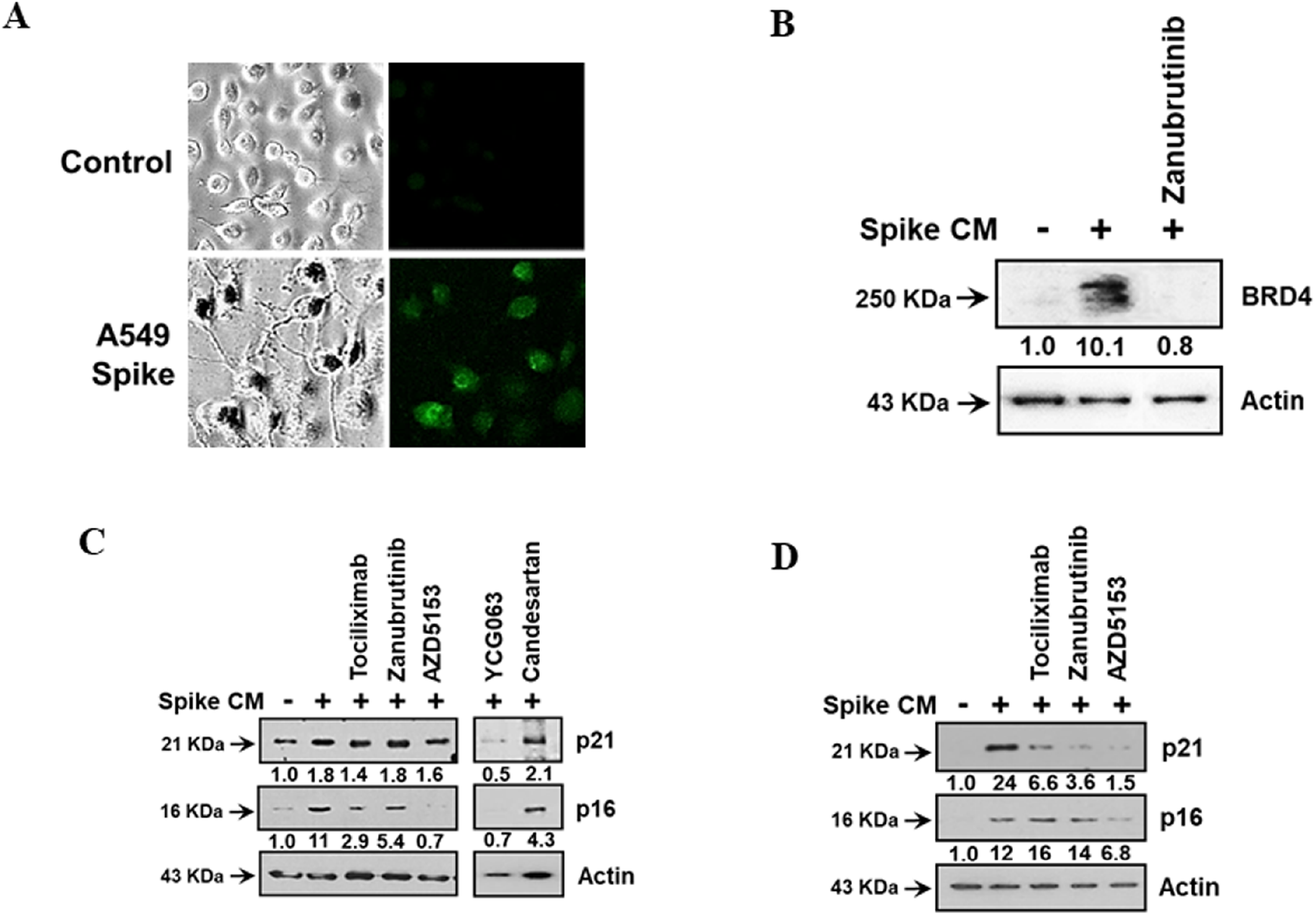
Cellular changes in TMNK by CM from SARS-CoV-2 spike expressing A549 cells. Culture medium from spike transfected A549 cells induced cellular changes in endothelial cell line TMNK and enhancement of SA-β-Gal (panel A). Exposure of TMNK-1 cells to A549 Spike expressing culture medium led to a marked induction of BrD4, which could be inhibited by Zanabrutinib (panel B). TMNK and EA_hy926_ cell lines exhibited enhanced expression of the senescence marker p16 and p21 by western blot (panels C and D). Experiments were performed in triplicate.

We further examined TMNK and EA cell lines for induction of the senescence related marker proteins, p16 and p21. Both proteins were induced in each cell line after exposure to A549 Spike conditioned media (Fig. 4, panels C and D). We had earlier identified an enhanced secretion of MCP1; a component secreted by the SASP, from TMNK cells treated with A549 Spike conditioned media, which could be inhibited by treatment with Tociliximab. Here, we treated these endothelial cell lines to inhibit cell response to external stimuli with a variety of inhibitors focused upon impeding cytokine response or SASP function. Tociliximab displayed the ability to impede p16 expression in TMNK cells, but had little effect upon reducing expression in EA cells.

We also used the BTK inhibitor; Zanabrutinib to inhibit cytokine response along these pathways. Zanabrutinib failed to inhibit p21 expression, but did reduce p16 expression. In EA_hy926_ cells, Zanabrutininb could reduce the level of p21, but not p16. We further analyzed the use of the Brd4 inhibitor; AZD5153, noting that p21 was not reduced, but p16 was significantly inhibited. In contrast, AZD 5153 could significantly reduce the expression of p21 and p16 in EA_hy926_ cells.

Additionally, the use of the ROS inhibitor (YCG063) led to a decrease in both p21 and p16; in treated TMNK cells; indicating a potential role for ROS accumulation in endothelial cell pathology. Interestingly, in agreement with our previous work, the use of the AT1R inhibitor Candesartan cilexetil (which modulated Il-6 release from Spike expressing A549) did not significantly inhibit the presentation of either senescence marker in TMNK cells.

Endothelial cells produce various cytokines and chemokines in response to inflammatory processes. Stimulation of endothelial cells with recombinant human IL-1β or TNF-α induced cell associated bioactive cytokine release (21). TMNK cells exposed to CM from Spike transfected A549 cells exhibited an increase in the expression of IL-1α, IL-6 and MCP-1; but did not show modulation of HMGB1 (Fig. 5). The use of Tociliximab, Zanubrutinib, or AZD5153 greatly reduced the secreted levels of IL-6 and MCP-1. These results indicated that soluble mediators released from Spike expressing A549 cells may induce a senescence phenotype (SASP) in surrounding endothelial cells, and that this response may be modulated by specific inhibitors.

**FIGURE 5.**
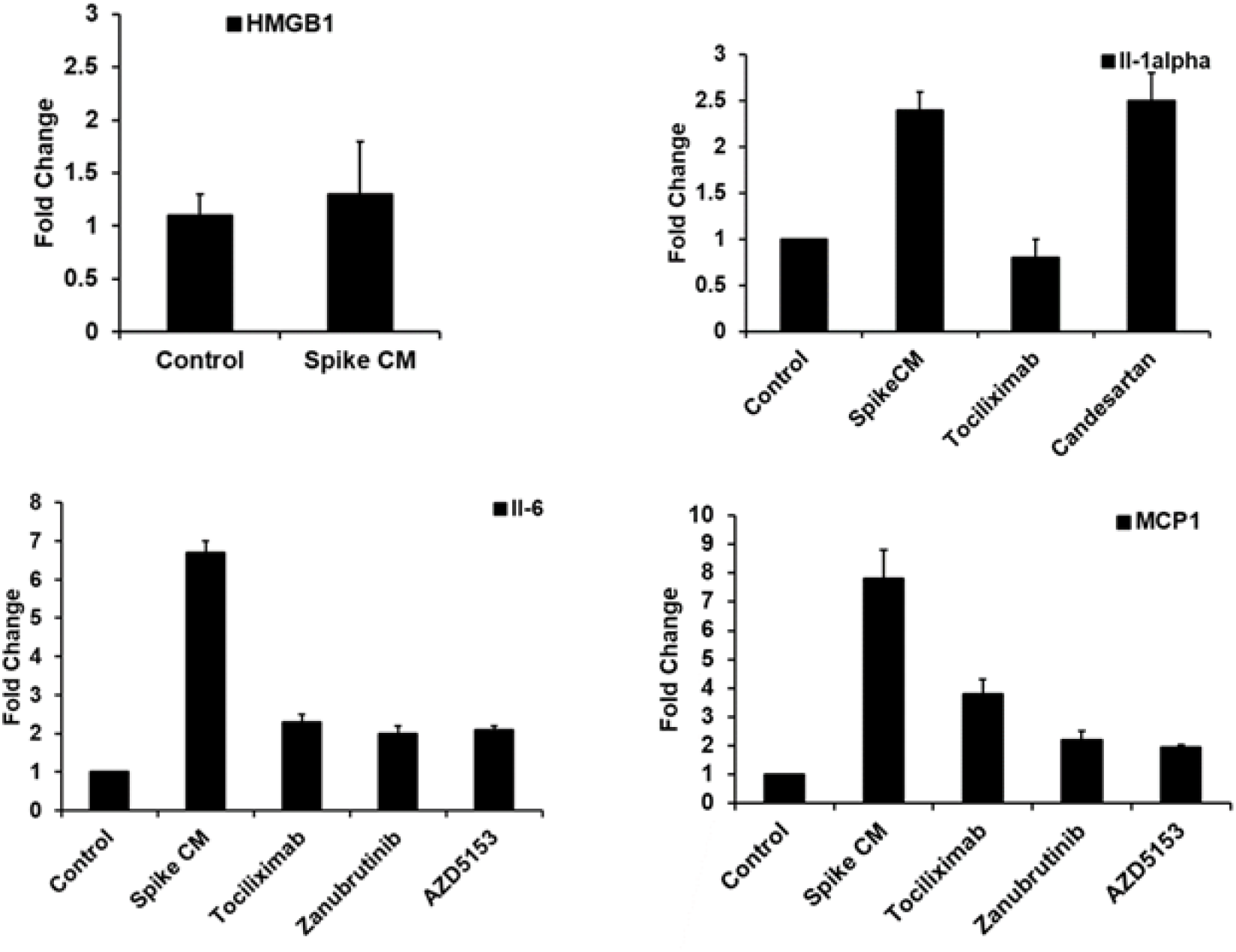
Release of cytokines and chemokine from TMNK cells following incubation with CM from SARS-CoV-2 spike expressing A549 cells. The resulting change in secretory cytokines and chemokine secretion for HGMB1, IL-1α, IL-6, and MCP-1 in the presence of inflammatory inhibitors are shown. Experiments were performed in triplicate.

### A549 Spike transfected culture media induces ROS generation in endothelial cells

Immune-pathogenesis of many diseases associate with enhanced ROS stress (22–24). Elevated levels of intracellular reactive oxygen species (ROS), are commonly observed in senescent cells (25). We examined ROS production in endothelial cells exposed to A549 Spike conditioned media in the presence of specific inhibitors. ROS levels were increased approximately 2-fold in endothelial cells (TMNK-1 and EAhy926) exposed to A549 Spike conditioned media, in comparison to those exposed to control media (Fig. 6, panels A and B). The use of YCG063 as a positive inhibitor control exhibited ROS levels below TMNK exposed to control media (data not shown). STAT3-related cytokines (i.e IL-6) activate Janus tyrosine kinases (JAK), triggering STAT3 phosphorylation and nuclear translocation, where it functions as a transcriptional factor and induces cellular senescence (26). IL-6 has been observed to enhance ROS levels in endothelial cells by upregulating AT1R gene expression. As shown in our previous work, endothelium does not express transmembrane IL-6Rα and is unresponsive to IL-6; however, endothelium can be activated by the IL-6 trans-signaling pathway. Here, use of Tociliximab, a known inhibitor of IL-6 signaling, inhibited ROS production in endothelial cells when cultured in the presence of A549 Spike CM.

**FIGURE 6.**
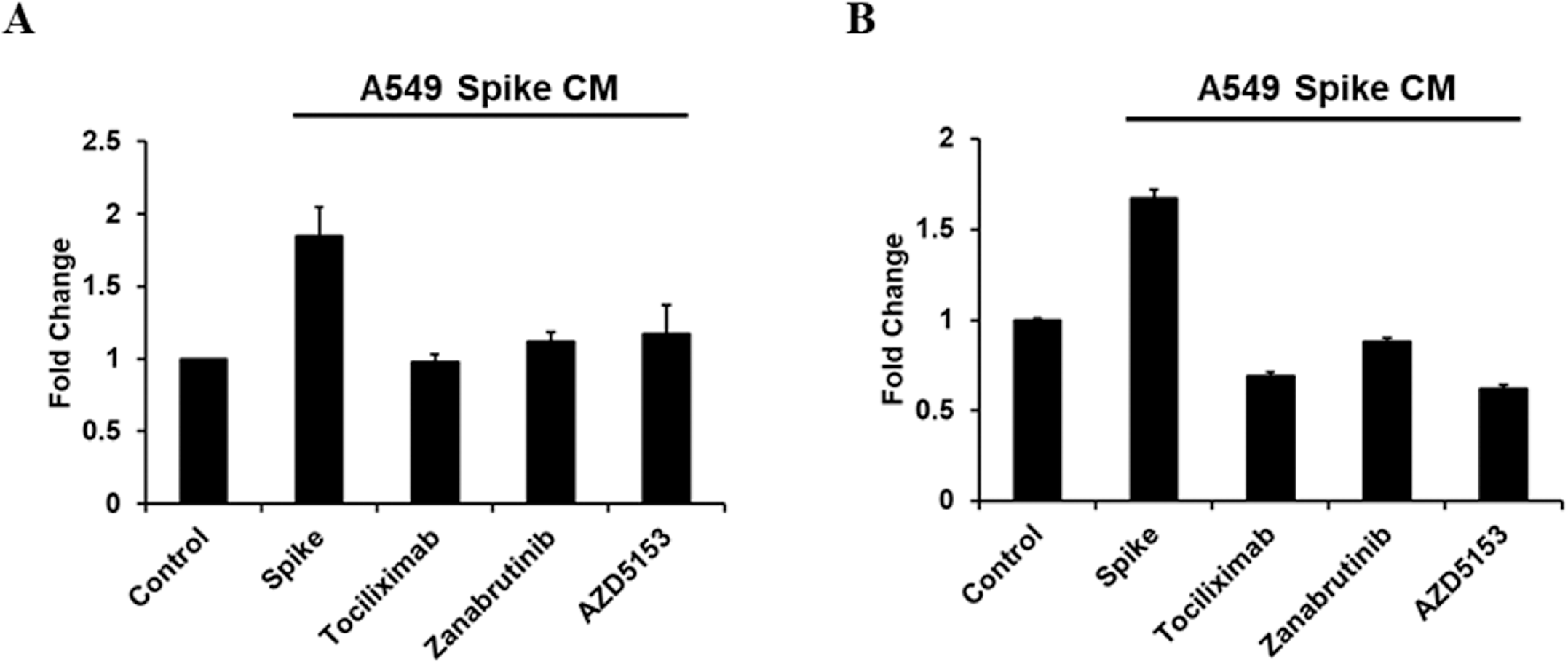
A549 Spike transfected culture media induces ROS generation in endothelial cells. ROS production in endothelial cells exposed to A549 Spike conditioned media in the presence of specific inhibitors. ROS levels were increased approximately 2-fold in endothelial cells exposed to A549 Spike conditioned media, in comparison to those exposed to control media (panels A and C Experiments were performed in triplicate.

We further tested inhibitors which reduced senescence marker expression, and found that each of these were also effective in reducing ROS under similar conditions, in both TMNK and EA_hy926_ cells. These results indicated that the production of ROS in SARS-CoV-2 spike CM exposed endothelial cells may be modulated by reducing reactivity to both IL-6, by reducing senescence associated reactivity of the MyD88 and BTK pathways, and by inhibiting SASP function.

Activated AKT expression promotes accumulation of p53 and p21, increases cell size and induces senescence-associated β-galactosidase activity. We analyzed Akt activation in endothelial cells exposed to A549 Spike CM, and noted a modest increase in Akt activation on TMNK cells, with a more significant effect upon EA_hy926_ cells. MYD88 signaling may induce activation of the p38 and Akt pathways via BTK and SYK (27, 28). Zanabrutinib decreased phosphorylated Akt expression in both TMNK and EA_hy926_ cells.

### Modulation of Akt and p38 activation by viral Spike expressing A549 conditioned medium

Senescent cells are characterized by increased ROS levels. p38, a senescence associated mitogen-activated protein kinases (MAPK) is activated by cellular stresses including ROS, UV-and γ-radiation, and proinflammatory cytokines (29). p38 plays an important causative role in cellular senescence attributed to oxidative stress (30). The inhibition of p38 activation in H_2_O_2_ derived senescent cells led to a modest reduction in SA-β-gal expression, and a decrease in ROS production (31). Here, p38 activation was observed in both endothelial cell lines 24 hours after exposure to A549 Spike CM, and this activation was decreased by treatment with Tociliximab, Zanabrutinib, or AZD5153 (Fig. 7, panels A and B). Inhibition of ROS by YCG063 or inhibition of AT1R by Candesartan ciletaxil upon TMNK cells had no effect upon inhibiting these two pathways.

**FIGURE 7.**
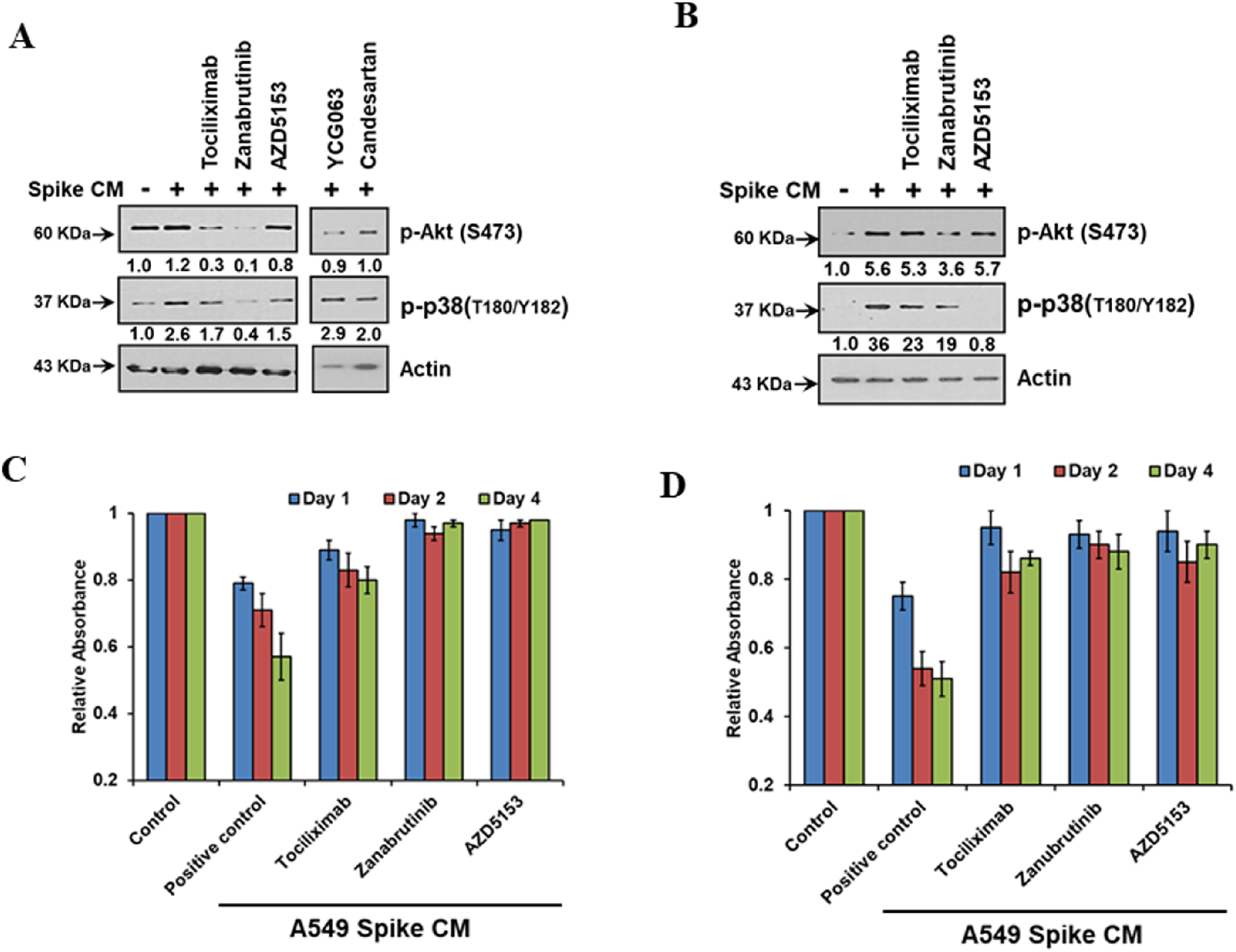
Modulation of Akt and p38 activation by viral Spike expressing A549 conditioned medium in endothelial cells and inhibition of cell proliferation. TMNK and EA_hy926_ cells were exposed to A549 Spike CM for 24 hours and measured for the phosphorylation of p38 and Akt by western blot. At exposure, individual wells were treated with specific inhibitors as indicated (panels A and B). **T**he proliferative capacity of endothelial cells treated with control or A549 Spike conditioned media (CM), and the effect of inhibitor treatment. A549 Spike CM decreased the proliferative ability of the endothelial cell line as displayed by progressively reduced MTT activity, as compared to control CM (panels C and D). Experiments were performed in triplicate. The results shown in panel D are from duplicate assays.

### Exposure of endothelial cells to A549 Spike transfected culture medium exhibits decreased proliferation

Senescence is an irreversible arrest of cell proliferation while the cell maintains metabolic function. Here, we compared the proliferative capacity of endothelial cells treated with control or A549 Spike conditioned media (CM), and the effect of inhibitor treatment. A549 Spike CM decreased the proliferative ability of the endothelial cell line as displayed by progressively reduced MTT activity (44% on TMNK and 48% on EAhy926 reduction apparent on day 4 post-exposure), as compared to control CM (Fig. 7, panels C and D). The use of Zanabrutinib (BTK inhibitor), or AZD5153 (Brd4 inhibitor) restored the proliferative capacity of the CM treated cells, with Tociliximab also able to limit the loss of proliferative capacity in treated cells.

### Senescent endothelial cells express adhesion molecules for leukocyte attachment

Adhesion molecules, vascular cell adhesion molecule 1 (VCAM1) and intercellular adhesion molecule 1 (ICAM1) are expressed on the surface of endothelial cells in response to cytokines, which contribute to leukocyte attachment for initiating vasculopathy or thrombosis. We observed that the endothelial cell lines, TMNK-1 and EA hy926, exhibit a senescence state after introduction of culture supernatant from SARS-CoV-2 Spike expressing epithelial cells. Enhanced ICAM-1 associated with endothelial cell senescence (32). We examined whether these senescent endothelial cells express enhanced adhesion molecules. Our western blot analysis showed that both TMNK-1 and EA hy926 cells express an increased amount of ICAM-1 and VCAM-1 molecules after exposure of culture supernatant from SARS-CoV-2 spike expressing A549 cells (Fig. 8, panels A and B). These enhanced ICAM-1 and VCAM-1 molecules were reduced after rescue from senescence state following treatment with Tocilizumab, Zanabrutinib, ST2825, AZD5153 (Fig. 8, panels A and B). SASP derived cytokines modulate leucocyte trafficking increasing adhesion molecule expression on epithelial cells, and through the induction of senescence in bystander cells (33). To determine further whether these contrasting phenotypes are also seen in more physiological conditions, we assessed leukocyte adhesion using a fluorometric based assay on the human monocyte cell line, THP-1. TNF-α has been observed to stimulate leucocyte -endothelial interactions via the upregulation and activation of cell adhesion molecules, and was included as a control (34). A significant cellular adhesion of THP-1 cells on the surface of TMNK-1 and EA hy926 cells were detected in the presence of culture supernatant from SARS-CoV-2 spike expressing A549 cells in association with elevated expression of the adhesion molecules (Fig. 8, panels C and D). The loss of leukocyte adhesion also corroborated with the reduction of ICAM-1 and VCAM-1 after treatment with Tocilizumab, Zanabrutinib, ST2825, AZD5153 (Fig. 8, panels C and D). These results indicate that leukocyte adhesion occurs to endothelial cells in a cellular environment, which is influenced by SARS-CoV-2 Spike expression in surrounding epithelial cells.

**FIGURE 8.**
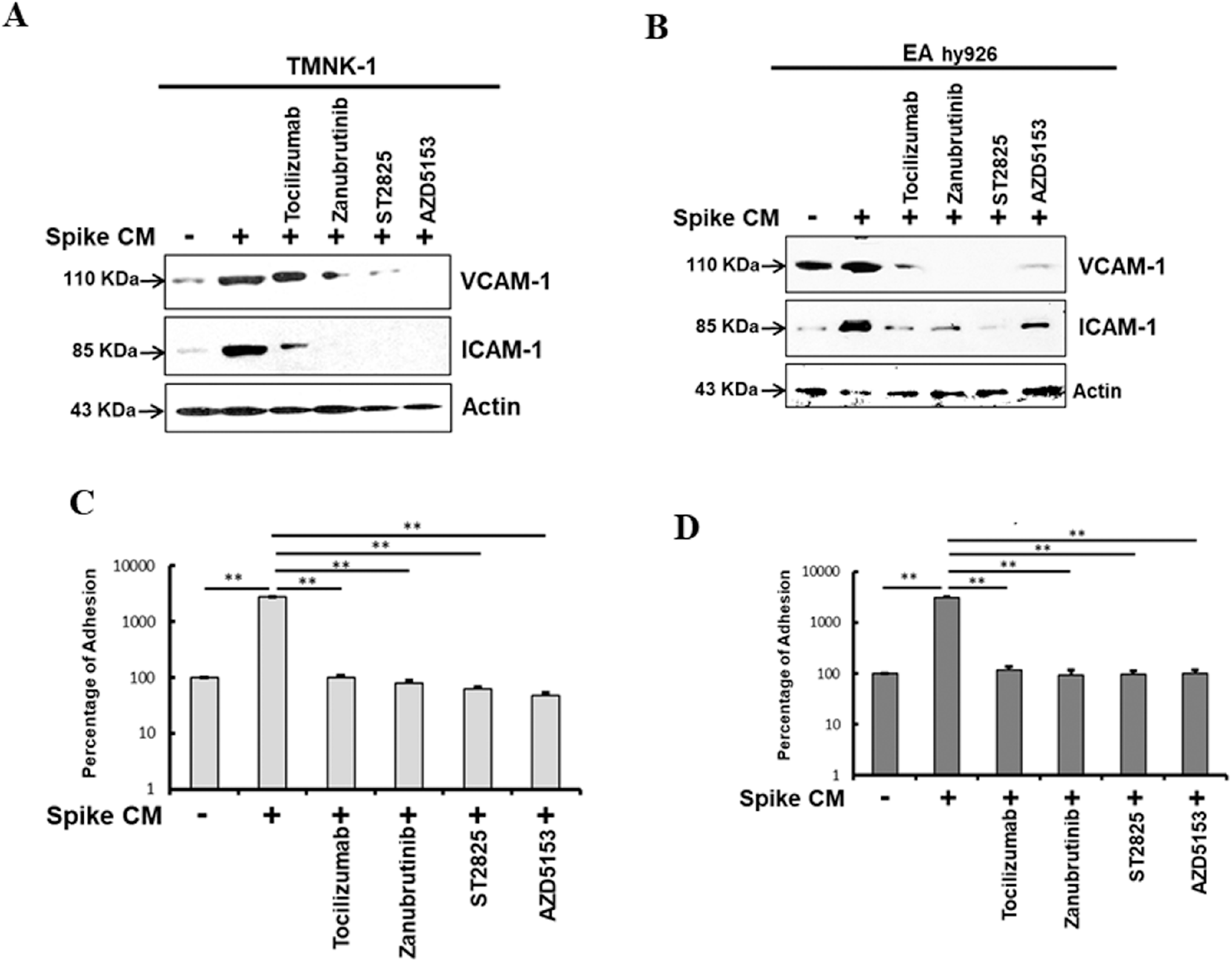
Bystander senescence induces endothelial adhesiveness. Expression of adhesion molecules VCAM-1 and ICAM-1 were analyzed by western blot from TMNK-1 and EA _hy926_ cell lysates prepared after 24 h exposure in culture supernatant of A549 cells in the presence or absence of SARS-CoV-2 spike protein with or without treatment of Tocilizumab, Zanubrutinib, ST2825 and AZD5153 (Panels A and B) Expression level of actin in each lane is shown as a total protein load control for comparison. A Comparative analysis of leukocyte adhesion on endothelial cells after 24 h exposure of culture supernatant of A549 cells in presence or absence of SARS-CoV-2 spike protein with or without treatment of Tocilizumab, Zanubrutinib, ST2825 and AZD5153 (Panels C and D). The percentage of adhesion was determined by a release of fluorescence from human monocytes (THP-1) after 6 h adhesion on treated TMNK-1 and EA hy926 cells. The results are presented as mean ± standard deviation. ‘*’ and ‘**’ represent statistical significance of p<0.05 and p<0.005, respectively. Experiments were performed in duplicate.

## Discussion

Our previous study revealed a marked elevation of IL-6, IL-1α and HMGB-1 in the culture supernatant from SARS-CoV-2 Spike expressing epithelial cells. These inflammatory molecules may be released through the senescence associated secretory pathway. Here, we have shown that SARS-CoV-2 Spike protein expression induces senescence in epithelial cells. SARS-CoV-2 Spike protein promotes IL-6/IL-6R induced trans-signaling and inflammatory responses that modulate MCP-1 expression in endothelial cells. These cells released alarmins and induced endothelial cell senescence. Spike transfected pulmonary epithelial cells (A549) exhibited an increase in senescence related p16 and p21 marker proteins and SA-β-galactosidase. Alveolar type-II lung cells harboring SARS-CoV-2 were shown to exhibit senescence with a proinflammatory phenotype in association with the presence of spike protein in COVID-19 patients (13).

Cellular senescence is an inducer of stress in tissues harboring senescent cells, leading to the expansion of senescence to normal bystander cells (35). The senescence bystander effect is initiated via NFκB mediated signaling induced by reactive oxygen species produced in senescent cells, and by the production of cytokines attributed to the function of the senescence associated secretory phenotype (SASP) (36). The stress response kinase p38 is a major contributor to the upregulation of NFκB (37), and the inhibition of p38 activation suppressed bystander senescence.

Indeed, activated p38 was observed in both SARS-CoV-2 infected and Spike transfected A549 cells. This finding corresponds with the prior result (38) which observed that the activation of NF-kB in both infected and Spike transfected A549 and Huh7.5 cell lines.

These observations lead to further examination of whether introduction of culture supernatant from SARS-CoV-2 Spike expressing epithelial cells generate senescence in endothelial cells. TMNK cells exposed to cell culture supernatant from A549 expressing SARS-CoV-2 Spike protein exhibited signs of senescence with enhanced p21, p16 and SA-β-galactosidase expression. Inhibition of IL-6 trans-signaling by Tocilizumab prior to exposure of supernatant to endothelial cells inhibited p16 and p21 induction, and reduced secretion of IL-1α. MCP-1 is secreted as a major component of senescence associated secretory phenotype (SASP). Senescence lead to an enhanced ICAM and VCAM expression in endothelial cells which may cause an increase in leukocyte adhesion with coronary blockade potential. Inhibition of alarmin associated signals or senescence regulation prevents ICAM/VCAM expression and leukocyte attachments. ROS was increased in Spike transfectants A549 and in TMNK cells exposed to transfected A549 culture supernatant. ROS generation in TMNK was reduced after treatment with specific inhibitors

Tociliximab, Zanabrutinib, AZD5153, and ST2825. Thus, our results indicated that the SASP related cytokines present in the culture supernatant from SARS-CoV-2 Spike expressing epithelial cells lead to senescence in endothelial cells. Blocking TLR or other inflammatory receptor mediated signaling prevented the paracrine effect in endothelial cells (Fig. 9).

**FIGURE 9.**
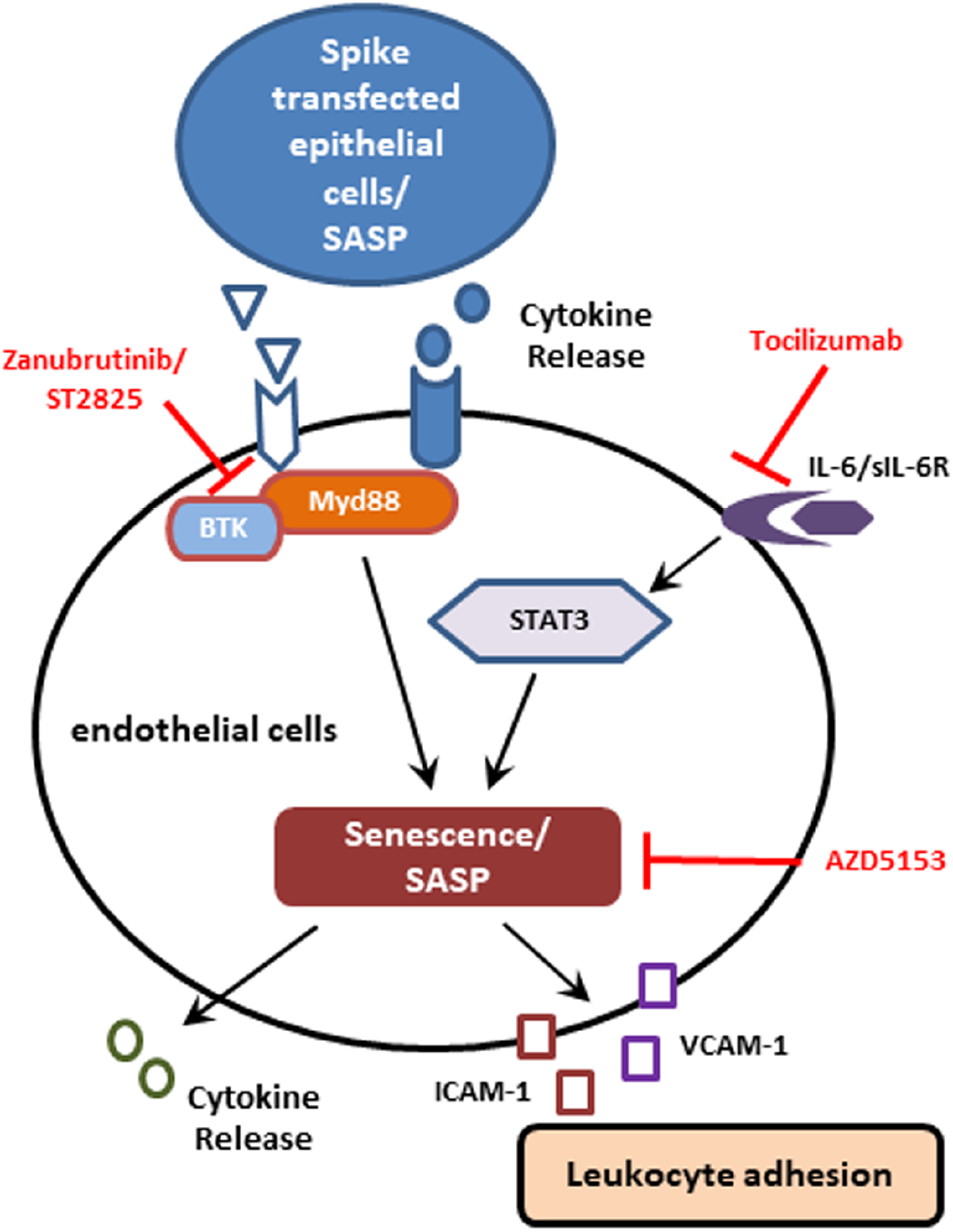
Schematic presentation of viral spike expressing epithelial cells leading to downstream signaling for functional consequences. Potential inhibitory steps of these signaling events are shown.

CD4^+^ and CD8^+^ T-cell counts are significantly reduced in COVID-19 patients, and the surviving T cells appear functionally exhausted (39). Patients with COVID-19 had increased CD8^+^ T cells expressing CD57, which is considered a key marker of *in vitro* replicative senescence and is associated either to human aging or prolonged chronic infections (40, 41).

ROS is produced as the byproduct of cellular metabolism. Accumulation of ROS causes cytostatic effects due to their ability to damage DNA, protein, and lipid molecules within the cell. ROS causes variety of lesions in DNA molecule which leads to DNA double strand breaks (42). Oxidative stress mediated DNA damage accelerates cellular senescence in the endothelial cells (43). Intracellular ROS production also leads to endothelial cell dysfunction and combine with NO to form peroxynitrite resulting in endothelial cell dysfunction (44).

Recent studies observed that Bromodomain-containing protein 4 (BRD4) is a novel regulator of the senescence associated secretory pathway. We observed that inhibition of BRD4 by AZD5153 reduces the release of SASP related cytokines, as well as senescence associated marker proteins. These SASP related cytokines function as paracrine mode to bind with their receptors located at membrane of the endothelial cells and promote pathogenesis. Our previous study also demonstrated that treatment with culture supernatant from SARS-CoV-2 Spike expressing epithelial cells containing elevated IL-6 resulted in activation of STAT3 tyrosine phosphorylation to induce MCP-1 expression in endothelial cells, and addition of Tocilizumab in culture supernatant inhibited MCP-1 expression. Taken together, we have identified that the exposure of endothelial cells to cell culture supernatant derived from SARS-CoV-2 Spike protein expression displayed cellular senescence markers.

Our study reveals that virus infected endothelial cells undergo senescence. The SASP release from the senescent cells serve as a paracrine signal to other tissues or cells, inducing senescence in distant organs or tissues. The reversal of senescence demonstrated in this study may be extended to clinical applications to alleviate disease severity and as a potential adjunct therapy to reduce COVID-19 mortality.

## Acknowledgments

We thank Daniel Hoft from Infectious Diseases, James Brien from Molecular Microbiology & Immunology, and Ratna B. Ray from Pathology in supporting our study.

## Conflict of interest

The authors declare that they do not have conflict of interest.

## Abbreviations used in this article

CM: culture media
ROS: reactive oxygen species
BRD4: bromodomain-containing protein 4
ICAM-1: intercellular adhesion molecule 1
VCAM-1: vascular cell adhesion molecule 1
SASP: senescence associated secretory phenotype
IL: interleukin
TNF: tumor necrosis factor
MCP-1: monocyte chemoattractant protein-1
SA-β-gal: senescence associated beta-galactosidase
DDR: DNA damage response
BTK: Bruton’s tyrosine kinase

## Notes

The study was supported by seed grant funding from Saint Louis University. The funders had no role in study design, data collection and analysis, decision to publish, or preparation of the manuscript.

### Competing Interest Statement

The authors have declared no competing interest.

